# Transcriptional Signatures of the Hierarchical Chronnectome Organization in the Human Brain

**DOI:** 10.1101/637942

**Authors:** Jin Liu, Mingrui Xia, Xindi Wang, Xuhong Liao, Yong He

**Affiliations:** State Key Laboratory of Cognitive Neuroscience and Learning, Beijing Normal University, Beijing, 100875, China; Beijing Key Laboratory of Brain Imaging and Connectomics, Beijing Normal University, Beijing, 100875, China; IDG/McGovern Institute for Brain Research, Beijing Normal University, Beijing, 100875, China; School of Systems Science, Beijing Normal University, Beijing, 100875, China

## Abstract

The chronnectome of the human brain represents the dynamics of functional organization in interacting regions, but its organizational principle and the underlying molecular mechanisms remain unclear. Using task-free fMRI and postmortem gene expression data, we conduct a transcriptome-chronnectome association study to investigate the spatial configurations of dynamic brain networks and their associations with transcriptional signatures. We reveal a spatial layout of network dynamics in the human brain chronnectome that reflects the cortical hierarchy and myelin content spanning from primary to transmodal areas. We further identify the transcriptional signature of this layout, with the top-related genes enriched for the ion channel and mitochondria terms. Moreover, the expression of these genes significantly predicts brain dynamics-behavior coupling. These findings highlight the hierarchical organizing principle and underlying molecular basis of the spatial configurations of dynamic brain networks, thereby contributing to our understanding of the associations among gene expression, network dynamics, and behaviors.

---

The human brain is a highly dynamic complex system with the hallmark of spontaneous fluctuations in neural activity over time. The emerging chronnectomics framework^1, 2^ together with advanced functional neuroimaging techniques (e.g., resting-state functional magnetic resonance imaging, rfMRI)^3^ demonstrates certain nonrandom characteristics of dynamic brain networks, such as time-varying connectivity strength^4, 5^ and modular reconfigurations^6^, as well as cyclical network transitions between states^4, 5, 7^, providing insights into our understanding of the dynamic organization of the functional network topologies to support various cognitive functions^6–10^. Here, we aim to explore a critical but underappreciated issue of the hierarchical ordering in the spatial organization of networked brain dynamics and the underlying molecular mechanism.

The cortical hierarchy spanning from primary sensorimotor to transmodal areas is a fundamental organizing principle of the human brain^11, 12^. Such a general principle has been observed both in the microstructural characteristics, such as intracortical myelin content^13, 14^ and the development sequence of subcortical white matter myelination^15^, and in the macroscopic properties of brain functions, such as functional connectivity features^16^ and the cycling transitions of metastates^7^. From an information processing perspective, the hierarchical organization of the brain allows for the efficient encoding and integration of parallel and series communications from sensation to cognition. However, whether the spatial layout of dynamic networks of the human brain follows the principle of hierarchical ordering remains unknown.

If the spatial configurations of brain network dynamics reflect the general cortical hierarchy, we speculate that a substantial molecular program could be conserved for coding this layout of the human chronnectome. Recently, the integration of whole-brain gene expression in the postmortem brain with neuroimaging *in vivo* provides unprecedented opportunities to bridge the gap between the underlying molecular mechanisms and brain networks^17–19^. For instance, two previous studies observed that the modular and hub architectures of the static functional networks during rest are associated with gene expression involving ion channel activity^18^ and oxidative metabolism/mitochondria^20^, respectively. During task states, the convergence of dynamic streams of functional networks is linked with the expression of synaptic long-term potentiation genes^21^. Notably, these previous studies did not provide evidence for the transcriptional signatures underlying the functional network dynamics that reflect the temporal organization of the spontaneous fluctuations of neural activity in the resting human brain^1, 22^. Clarifying this issue will not only provide insight into the functional and biological correlates of the brain network dynamics but also have indispensable implications for the understanding and interpretation of chronnectomics in normal development, aging, and disorders. Moreover, if the chronnectome architectures are promoted by specific transcriptional signatures, we anticipate that the expression patterns of chronnectome-related genes would be able to predict the involvement of network dynamics in a variety of cognitions and behaviors, thereby establishing a direct association among gene expression, brain dynamics, and behavioral phenotypes.

To address these issues, we conducted a transcriptome-chronnectome association study to investigate the spatial configurations and underlying transcriptional profiles of brain network dynamics by employing the rfMRI data from Human Connectome Project (HCP; 681 participants)^23^ and the microarray-based gene expression data from the Allen Institute for Brain Science (AIBS; 1791 samples from the brains of six donors)^17^. We identified three categories of dynamic brain hubs in terms of their time-varying patterns in global connectivity and modular switching (See *Methods*) that characterize the cortical hierarchical organization from primary to transmodal areas. Then, we demonstrated that the patterns of network dynamics can be explained by the expression profiles of genes associated with potassium ion channels and mitochondria. Moreover, the expression pattern of the top-related genes significantly predicted the relationship between network dynamics and cognitive and behavioral phenotypes.

## Results

### Spatial pattern of the human chronnectome reflects cortical hierarchy

Based on the rfMRI data from the HCP dataset, we constructed individual dynamic brain networks with a multimodal 360-region parcellation^24^ using a sliding window-based approach^4–6, 8, 9^ and a framewise-based conditional correlation approach^25^, respectively (see “Effect of the Dynamic Network Construction Approach” for more details). To characterize the spatial organization of dynamic brain networks, two measures were utilized, including the temporal global variability (tGV)^10^ and the temporal modular variability (tMV)^6^. For a given brain node, the tGV measures the overall time-varying fluctuations of the connections linked to the node, while the tMV measures its modular switching across time (see *Methods* and *SI* for details). As a toy example, both tGV and tMV were jointly used to classify network nodes into four distinct categories (Fig. 1): shaker hubs with large global variations but limited module switching (↑tGV and ↓tMV, yellow), bi-active hubs with high dynamics at both the global and modular levels (↑tGV and ↑tMV, red), switcher hubs with frequent transitions between modules but lower global variations (↓tGV and ↑tMV, green), and stabilizer non-hubs with relatively stable temporal variations (↓tGV and ↓tMV, blue).

**Fig. 1.**
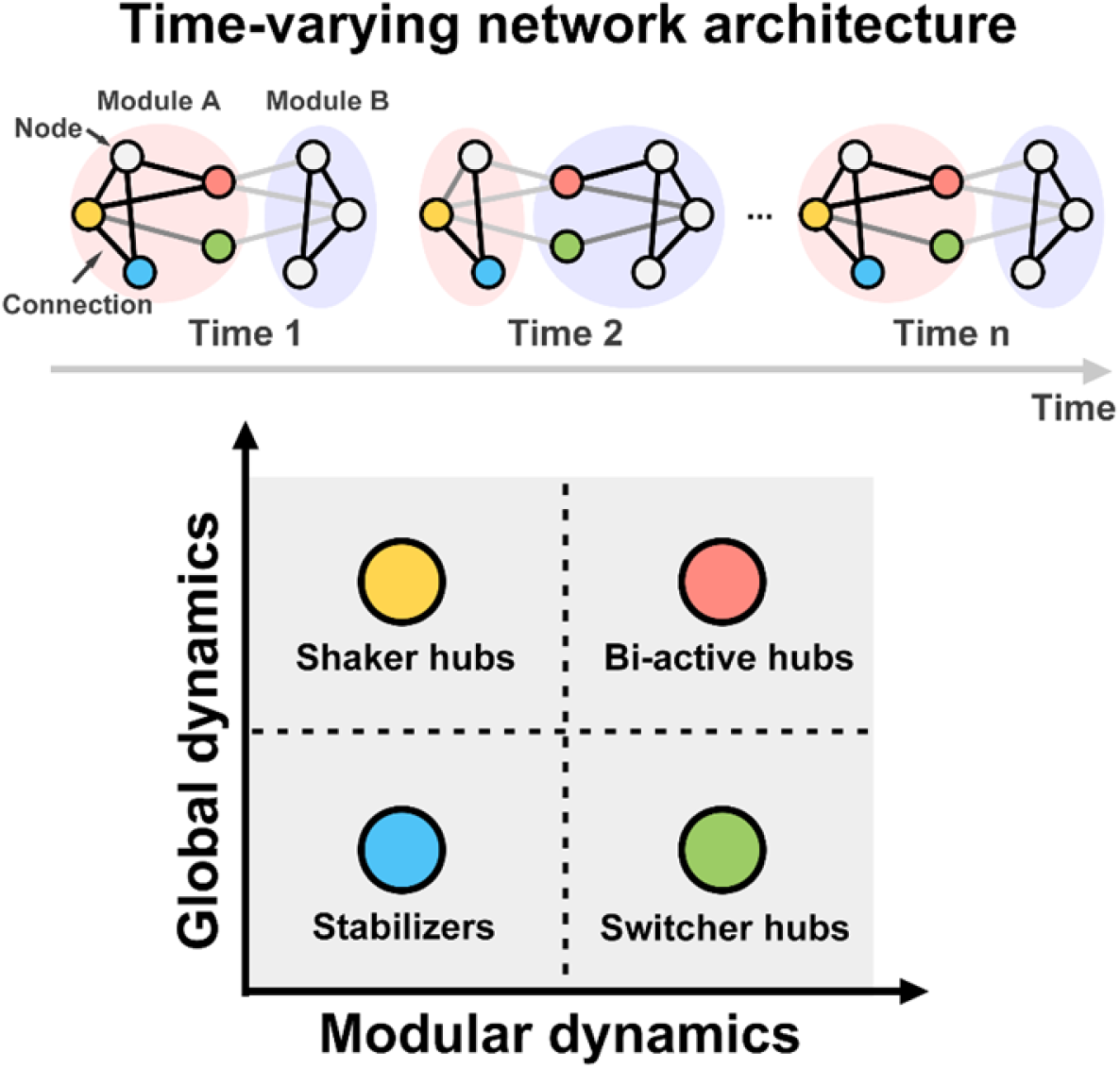
The toy model of different categories of network node dynamics. Different nodes have distinct patterns of temporal variations in global connectivity and modular affiliations across time. The strength of network connections from weak to strong is rendered from light gray to black. Each light circle represents a unique module composed of a set of nodes. The four categories of nodes are in different colors according to their dynamic metrics compared to the mean values of each measure: the shaker hubs exhibit large global fluctuations but limited modular dynamics (yellow); the bi-active hubs have notable global and modular dynamics (red); the switcher hubs show frequent modular variations but less global dynamics (green); and the stabilizer non-hubs exhibit very little global and modular dynamics (blue).

For each individual, we mapped their dynamic network activities in terms of the tGV and tMV maps. Both maps exhibited a high spatial similarity between sessions (tGV: *r* = 0.99, tMV: *r* = 0.97, all *p* < 0.0001) (Fig. S1A). The group-based, session-averaged tGV and tMV maps are illustrated in Fig. 2A and showed high similarity between the two hemispheres (tGV: *r* = 0.95, tMV: *r* = 0.97, all *p* < 0.0001) (see *SI*, Fig. S2, Fig. S3 and Fig. S4 for more details). We identified the three types of dynamic brain hubs (above brain mean in tGV, tMV or both) and non-hubs as follows (Fig. 2B): the shaker hubs were primarily in the sensorimotor cortex, involving the primary visual, motor, somatosensory and auditory cortices; the bi-active hubs were primarily in the lateral and medial frontal and parietal cortices; the switcher hubs were mainly in the anterior and middle cingulate cortices, medial temporal lobe, and anterior insula; and the stabilizer non-hubs were mainly in the sensorimotor association cortices, supplementary motor cortices, and posterior insula.

**Fig. 2.**
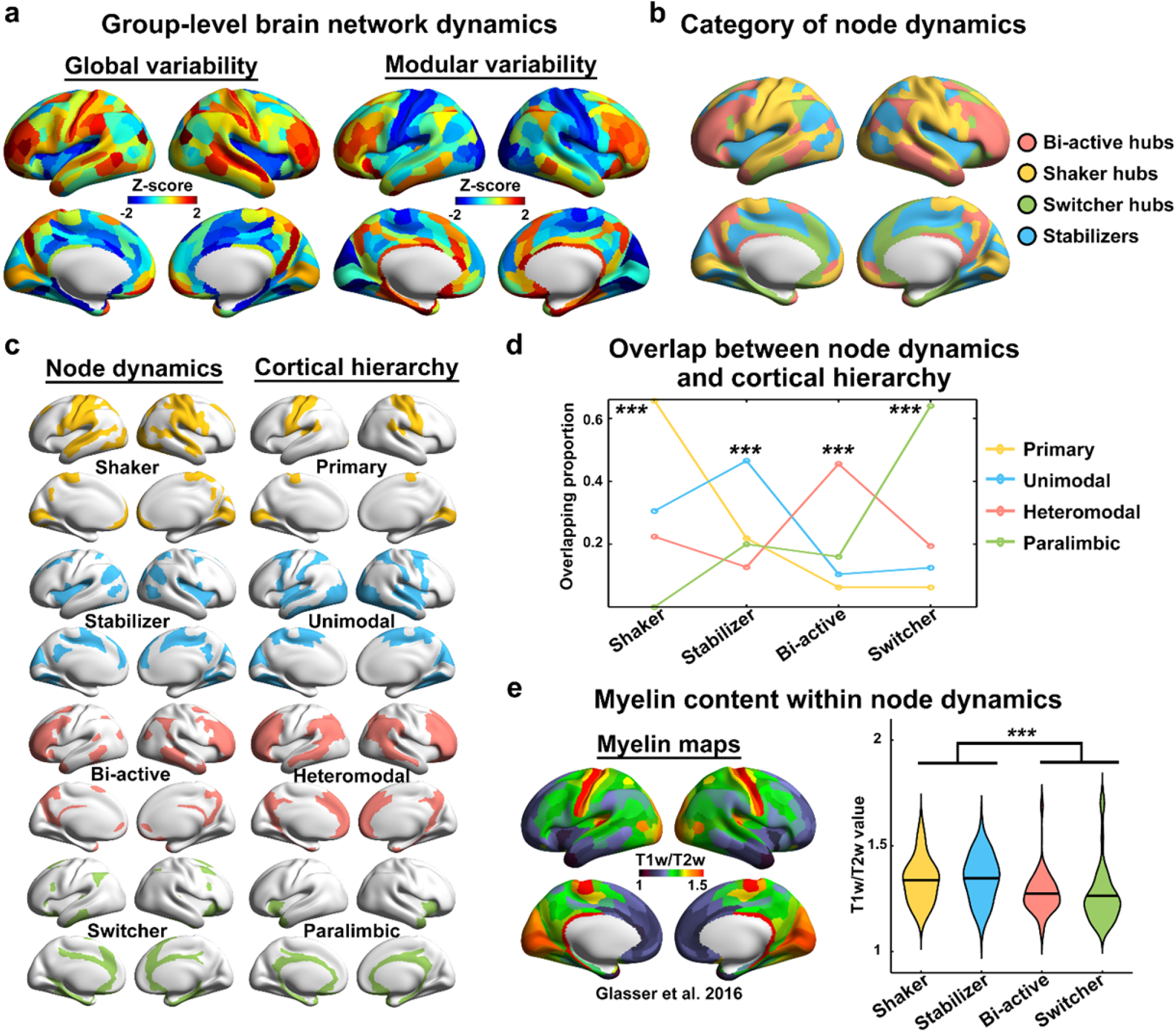
Spatial layout and microstructural relevance of the four categories of node regions in the dynamic brain networks. **a** The group-level maps of global and modular dynamics were obtained by averaging the individual brain activity maps. **b** The spatial distribution of the four categories of nodes in the dynamic brain networks. **c** The spatial distribution of each category of the dynamic network nodes (left column) and of each type of cortical hierarchy area (right column). **d** The overlap proportions of the four categories of network nodes in each type of cortical hierarchy area. Permutation tests were conducted to determine whether the proportions were significantly higher than those by chance. **e** Between-group differences in myelin contents across four categories of node dynamics. The myelin map estimated by the ratio of T1/T2-weighted MRI was obtained from Glasser et al.^24^ (left). The bi-active hubs and switcher hubs were significantly more lightly myelinated than the shaker hubs and stabilizer non-hubs (right, permutation test). *** *p* < 0.001. The surface rendering was using BrainNet Viewer (http://www.nitrc.org/projects/bnv/)^73^ with the inflated cortical 32K surface^24^.

To further explore whether the spatial configuration of brain network dynamics reflects the hierarchical organization, we assigned each node region to one of four types in the cortical synaptic hierarchy described by Mesulam^11^ (i.e., primary, unimodal association, heteromodal association, and paralimbic/limbic areas) (*Table S3*) and found that the spatial layouts of the four categories of dynamic network nodes were generally similar to the cortical hierarchy distributions (Fig. 2C and Fig. S1C). Furthermore, we calculated the overlap proportions of the four categories of nodes within each hierarchy and performed permutation tests to determine whether the proportions were larger than those by chance. We found that brain regions within each hierarchy were significantly dominated by a particular category of dynamic nodes (all *p* < 0.0001, 10,000 permutations): the primary area was mostly occupied by the shaker hubs, the unimodal association area was largely occupied by the stabilizer non-hubs, the heteromodal association area was primarily occupied by the bi-active hubs, and the paralimbic/limbic area was mainly dominated by the switcher hubs (Fig. 2D). Moreover, we compared the myelin content among the four types of dynamic nodes by employing a publicly available myelin map derived from the ratio of T1- and T2-weighted MRI^14, 24^, which has been considered a reliable noninvasive neuroimaging measure indexing cortical hierarchy^13^. We found that the transmodal hubs (i.e., bi-active and switcher hubs) exhibited significantly lighter myelination than the shaker hubs and stabilizers (all *p* < 0.0001, 10,000 permutations) (Fig. 2E). These findings suggest that the spatial configuration of the intrinsic chronnectome reflects a core synaptic hierarchy and is coupled with the cortical myeloarchitecture.

### Transcriptional signatures underlying the intrinsic chronnectome

To elucidate the molecular mechanism underlying the spatial layout of brain network dynamics, we explored the relationship between dynamic measurements and gene expression profiles of the adult human brain from the AIBS. Here, partial least squares regression^26^ was used to define components, each of which represents a linear combination of the gene expression in predictor variables (i.e., 16,392 genes) that can explain the variance in response variables (i.e., tGV and tMV).

We observed that two significant components from partial least squares regression explained a total of 28% of the variance in the dynamic measurements (*p* < 0.0001, permutation test, see *SI* and Fig. S5). Specifically, the first component represented an association between brain network dynamics and a transcriptional profile characterized by high expression mainly in the posterior parietal-occipital areas (Fig. 3A, left). The regional mapping of this component correlated positively with tGV but negatively with tMV. The second component represented an association between network dynamics and a transcriptional profile with high expression predominantly in the anterior prefrontal and temporal areas (Fig. 3A, right), the regional mapping of which correlated positively with both tGV and tMV. Finally, the transcriptional profile of the first component was significantly enriched in genes relating to the potassium ion channel complex and activity (all *q* < 0.001), and that of the second component was significantly enriched for a set of genes associated with the mitochondrial part of the cellular component (all *q* < 0.0001) (Fig. 3B, Fig. S6 and Fig. S7, *Table* S5). These findings were reproducible in the internal replication analysis (*Methods* and *Table S6*).

**Fig. 3.**
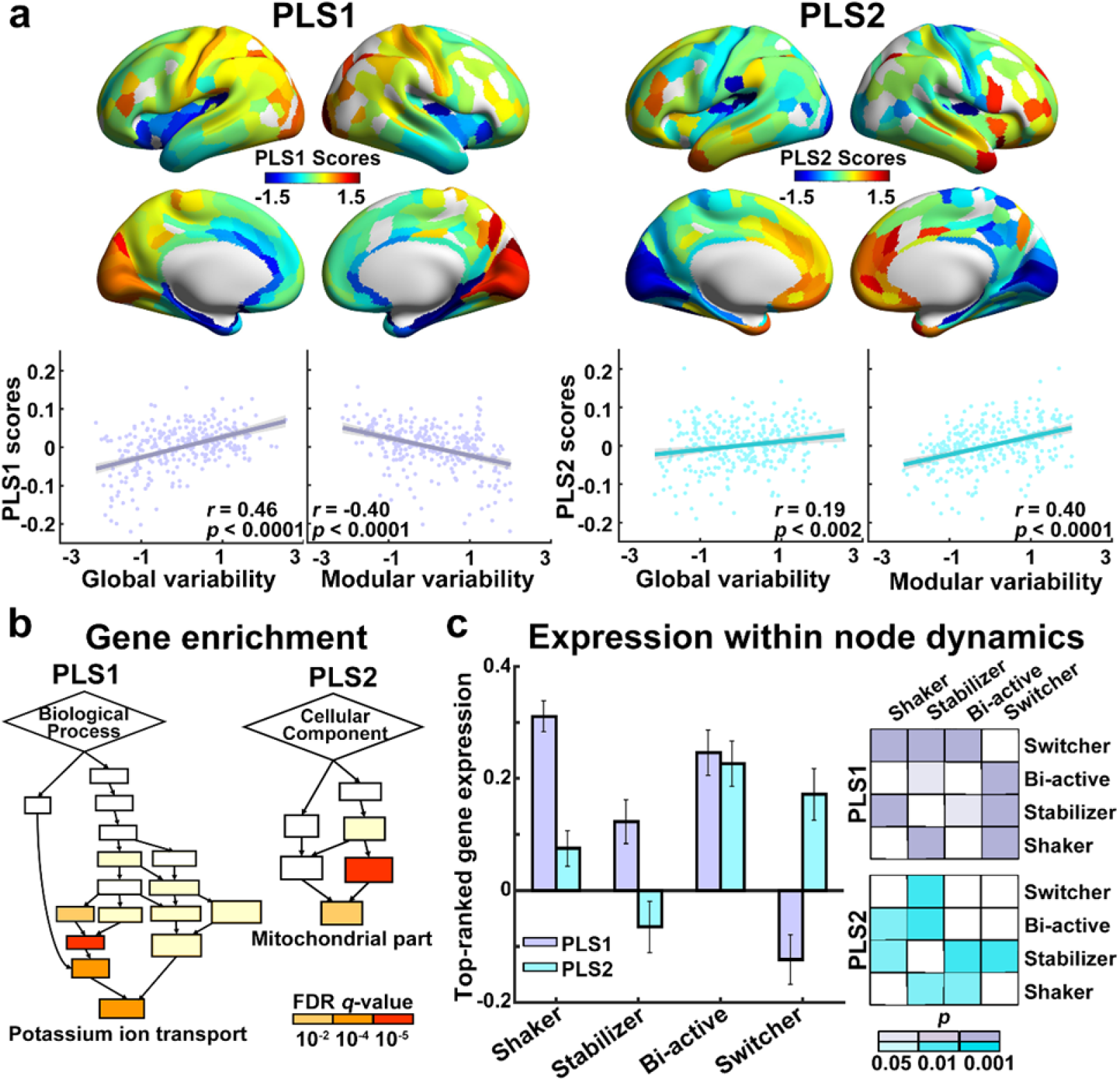
Association between gene expression profiles and dynamic network architectures. **a** The first two partial least squares (PLS) regression components (PLS 1 and PLS 2) explain 28% of the variance in the dynamic measures. PLS 1 identified a gene-expression profile with overexpression mainly in the medial occipital, lateral parietal, and lateral prefrontal cortices positively correlated with global dynamics but negatively correlated with modular dynamics (left column). PLS 2 identified a gene expression profile with overexpression dominantly in the medial/lateral prefrontal, and lateral temporal cortices positively correlated with both global and modular dynamics (right column). The shadow indicates the 95% confidence intervals. **b** PLS 1 is enriched for genes related to Gene Ontology of biological processes and cellular components associated with the complex and activity of potassium ion channels, whereas PLS 2 is enriched for genes related to the mitochondrial part (colors index the *q*-values for significant enrichment). See Fig. S5 and S6 in *SI* for more details about these Gene Ontology terms. **c** The mean expression levels of the top 10% weighted genes in PLS 1 and PLS 2 are diversely distributed across the four types of dynamic network nodes (left). The error bars indicate the standard error. There are significant between-group differences in the gene expression levels across the four types of nodes (right, colors index the *p*-values for significant between-group differences).

Significant differences in gene expression were observed among the four types of brain nodes (10,000 permutations, Fig. 3C). Specifically, for genes related to the potassium ion channel, higher expression was observed in the bi-active and shaker hubs than in the switcher hubs and stabilizers (all *p* < 0.018) and in the stabilizers than in the switcher hubs (*p* < 0.0001). For genes associated with mitochondria, higher expression was observed in the bi-active hubs than in the shaker hubs and stabilizers (both *p* < 0.004) and in the switcher hubs and shaker hubs than in the stabilizers (both *p* < 0.009). These findings indicate specific transcriptional signatures underlying the different temporal dynamics of brain nodes in the chronnectome.

### Transcriptional signatures support the involvement of the intrinsic chronnectome in behaviors

To determine whether the transcriptional signatures of the intrinsic chronnectome significantly predict the regional involvement of network dynamics in cognition and behavior, we used a machine-learning method based on support vector regression^27^. Specifically, we estimated the regional behavioral involvements by implementing a canonical correlation analysis (CCA)^28^ onto the brain network dynamics and 92 behavioral measures from the HCP (*Table S1).* The CCA can identify the modes that relate two data sets by identifying the optimal linear combination in each with the maximal correlation. With the expression of top-ranked genes (10%) from two significant components of partial least squares regression as features, we performed support vector regression analysis to build models to predict the behavioral involvement measured as the total absolute weight of CCA mode on two dynamic measurements.

Our analysis revealed a significant CCA mode that related the intrinsic dynamic architectures with a set of behavioral measures (r = 0.79, *p* = 0.009, 10,000 permutations). The behavioral measures correlated with this CCA mode were primarily organized into a positive-negative axis. Specifically, the positively correlated behavioral measures mostly belonged to cognition or positive personal qualities, such as reading decoding and psychological well-being, while the negatively correlated behavioral measures largely overlapped with negative personal traits or problems, such as depressive and avoidant personality problems (Fig. 4A). Fig. 4B shows the regional weight of this CCA mode on two dynamic measures. Notably, the significant weights of the CCA mode were diversely distributed across the four categories of dynamic brain nodes. The bi-active hubs and the switcher hubs had significantly larger weights to the CCA mode than did the shaker hubs and stabilizers (all *p* < 0.006; Fig. 4C). Moreover, we validated this CCA mode with different parameters, with little change in the results (*Table S7*).

**Fig. 4.**
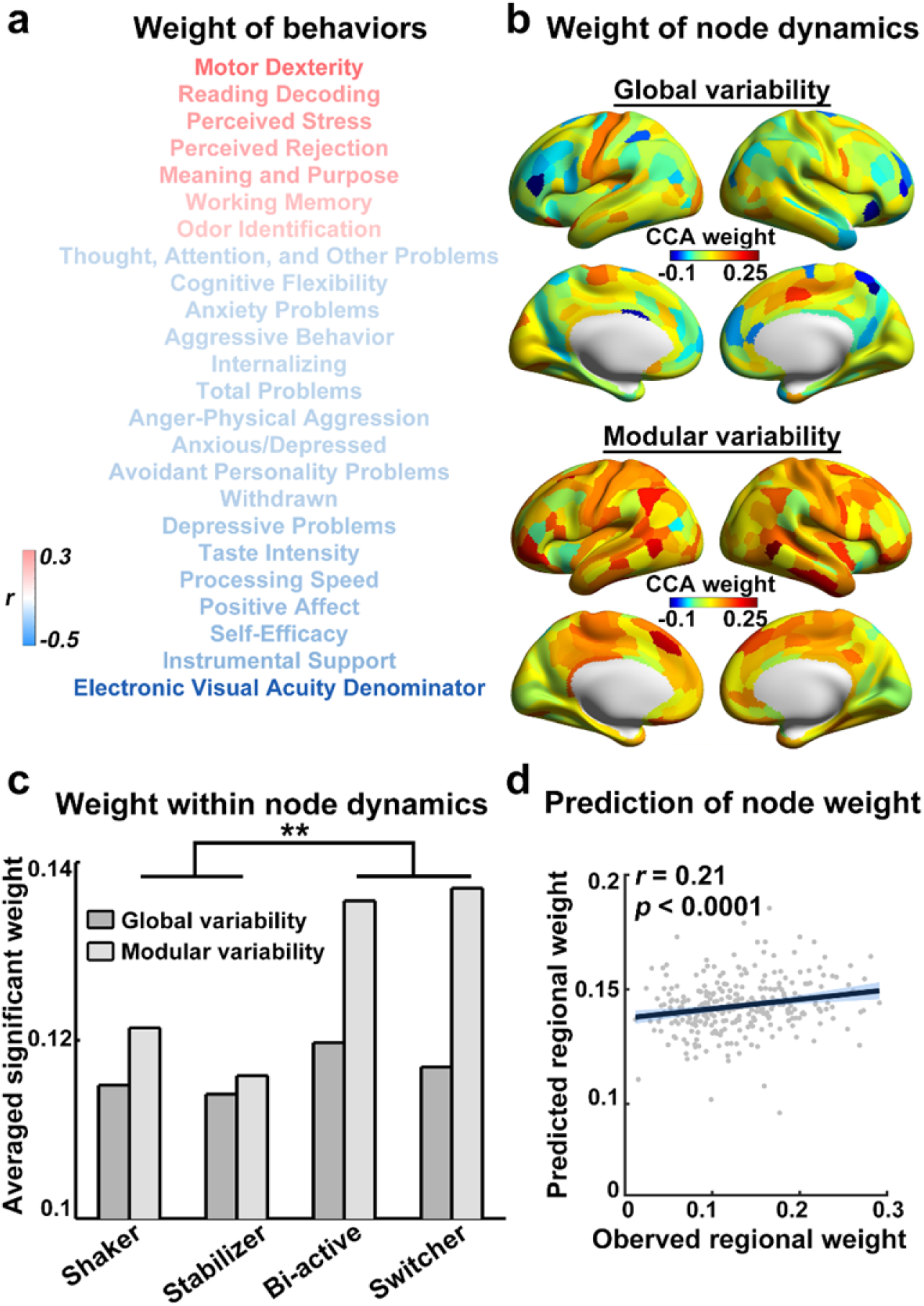
Association between gene expression and behavioral involvement of dynamic network architectures. **a** The CCA mode reveals that a set of individual behavioral measures is significantly correlated with global and modular brain dynamics, with positive correlations mostly associated with the cognition or positive personal qualities (red) and negative correlations mainly related to negative personal traits or problems (blue). **b** The spatial distribution of the regional weight of the CCA mode on two dynamic measures. **c** The weights of the CCA mode for significant brain-behavior correlations (*p* < 0.01) on two dynamic measures were diversely distributed across the four types of dynamic network nodes, and between-group differences were identified using permutation tests. **, *p* < 0.01. **d** Scatter plot shows the correlation between the predicted and observed regional cognitive involvement (sum of the absolute CCA mode weight on two dynamic measures). The prediction was performed using the support vector regression analysis with chronnectome-related gene expressions as features. Each dot represents the data of one node, and the shadow indicates the 95% confidence intervals.

We found that the expression patterns of chronnectome-related genes significantly predicted the weight of network dynamics in the brain CCA mode. Based on a feature-selection threshold of *p*-value < 0.01, the correlation between the observed CCA weight and the predicted CCA weight was *r* = 0.21 (*p* < 0.0001, 10,000 permutations) (Fig. 4D). Based on another feature-selection threshold of p-value < 0.001, the prediction remained significant (r = 0.17, *p* < 0.0001) (Fig. S8). These findings suggest that these gene expression patterns may regulate the involvement of the intrinsic chronnectome architectures in the cognitive and behavioral phenotypes.

### Effect of head motion

To further evaluate the effect of head motion on our main findings, we performed spike-regression-based scrubbing in the nuisance regression procedure during imaging data preprocessing^29, 30^. We found that the spatial time-varying patterns were highly similar to our main results (tGV and tMV, both *r* > 0.99, *p* < 0.0001, permutation tests; node classification, NMI = 0.93) (Fig. S9A). The spatial patterns of network node dynamics corresponded well to the cortical hierarchy: the primary area was mainly occupied by shaker hubs, the unimodal association area was largely occupied by stabilizers, the heteromodal association area was mainly occupied by bi-active hubs, and the paralimbic/limbic area was mainly dominated by switcher hubs (Fig. S9B, left). The transmodal hubs (i.e., the bi-active and switcher hubs) were significantly less myelinated than the shaker hubs and stabilizers (*p* < 0.0001, Fig. S9B, right). Regarding the transcriptional signatures, two significant partial least squares regression components explained 30% of the variance in dynamic measurements (*p* < 0.0001); the related genes were enriched for Gene Ontology terms associated with the complex and activity of potassium ion channels and the mitochondrial part (*Table S8).* There were significant differences in the expression of top-ranked genes among the four types of dynamic network nodes, with the bi-active hubs exhibiting the highest expression (Fig. S9C). Moreover, we found that the expression pattern of the brain dynamics-related genes significantly predicted the weight of brain-behavior coupling in the CCA mode (r = 0.15, *p* < 0.021, Fig. S9D).

### Effect of the dynamic network construction approach

Based on the dynamic networks constructed by the framewise conditional correlation approach without considering sliding windows^25^, we obtained similar spatial dynamic patterns compared with our main results for the tGV map (*r* = 0.83, *p* < 0.0001), the tMV map (*r* = 0.87, *p* < 0.0001) and node classification layout (NMI = 0.42) (Fig. S10A). The distribution of the four categories of node dynamics mainly followed a cortical hierarchy, with the primary area mostly occupied by shaker hubs and stabilizer non-hubs, the unimodal association area largely occupied by the stabilizers, the heteromodal association area mainly occupied by the bi-active hubs, and the paralimbic/limbic area mainly dominated by the switcher hubs (Fig. S10B, left). Consistent with the main results, the transmodal hubs (i.e., the bi-active and switcher hubs) exhibited significantly lighter myelination than the shaker hubs and stabilizers (all *p* < 0.0002, Fig. S10B, right). Furthermore, the partial least squares regression analysis revealed three significant components explaining 38% of the variation in the dynamic measurements (*p* < 0.0001), and these components were partly associated with genes related to the ion channel or mitochondrial cellular component (*Table S9*). Significant differences in gene expression were observed among the four types of brain nodes, with the bi-active hubs having the highest expression (Fig. S10C). Finally, the expression patterns of chronnectome-related genes were able to significantly predict the weight of associations between network dynamics and behavioral measures in the significant CCA mode (*r* = 0.41, *p* < 0.0001, Fig. S10D).

## Discussion

In this study, we demonstrate the spatial heterogeneity of brain network dynamics in terms of temporal variations of global and modular organization. This architecture reflects hierarchical function processing and couples with the underlying myeloarchitecture. Importantly, the network dynamics are supported by substantial gene expression profiles, with brain nodes having diverse transcriptional signatures in the expression of genes regulating potassium ion channel activity and mitochondria. Moreover, the expression pattern of these genes significantly predicts regional functional involvement in a variety of cognitive and behavioral phenotypes. Collectively, these findings provide empirical evidence for the hierarchically organized architecture of the chronnectome and its neurobiological substrate and have implications for the understanding and interpretation of the organizational principle and working mechanism of brain network dynamics.

### The intrinsic chronnectome architectures reflect hierarchical function processing

We characterized the chronnectome architectures that are composed of three categories of dynamic brain hubs (i.e., the bi-active hubs, the switcher hubs, and the shaker hubs) and one category of non-hubs. The spatial distributions of these nodes with distinct time-varying features are highly comparable to the core synaptic hierarchy for information processing from sensation to cognition^11^ and the underlying myeloarchitectural features of the human brain^14^. Specifically, the shaker hubs with high global fluctuations are mainly located in the primary sensorimotor cortices that provide obligatory portals to access sensory stimuli from the external world. Thus, the function of primary areas addresses the most primary but continuous processing for particularly specialized sensory information with abundant synaptic connections. Such processing requires dramatically dynamic connectivity fluctuation for the constant monitoring and capturing of information from the ever-changing external environment. The stabilizers are predominantly located in the unimodal association areas that encode the basic features of sensation captured from particular sensory modalities, such as color, motion, and sound from the primary cortex, into complex contents of sensory experience, such as objects, faces, and spatial locations^11^. Given the nature of highly specialized modalities, these unimodal association areas with limited global fluctuations and intermodular switching fit this unique role in information processing. Interestingly, these two types of dynamic nodes belong to heavily myelinated areas, which have dense and well-organized neurons with small dendritic arbors and are myelinated earlier during development^15, 31^. Thus, the relatively simple myeloarchitectural features of the shaker hubs and the stabilizers are in line with their functional roles in processing unimodal content reflected by less modular switching.

Importantly, the bi-active hubs with both high global and modular dynamics are mainly distributed in heteromodal association regions (i.e., the lateral and medial frontal and parietal cortices). These regions play central roles in forming distributed but integrated multimodal representations and are associated with high areal expansion during development and evolution^32^. The participation of the heteromodal association areas in receiving convergent inputs from multiple unimodal areas and binding with other transmodal areas usually requires global communications across distributed regions and integration among functional systems^33, 34^. Thus, the prodigious global connectivity fluctuations and frequent modular switching might provide a neurobiological foundation that enables the bi-active hubs to meet the high requirement of complex processes in high-order cognition in the human brain. The switcher hubs with only high modular reorganization are predominantly located in the limbic and paralimbic areas. These regions are related to the internal milieu and are responsible for the regulation of emotion and motivation. The literature suggests that the limbic and paralimbic areas are a complex organization responding to the interconnection between the primitive subcortical and the evolved cortical structures^35^. Both the bi-active hubs and switcher hubs belong to the transmodal cortices and are located in the lightly myelinated areas with low neuron density but complex dendritic arbors^36^. These observations are compatible with our findings that the two types of dynamic hubs are mostly involved in sets of behaviors. Therefore, the high temporal variations of modular structure in the bi-active hubs and the switcher hubs might provide a network foundation for the dynamic combination and integration of multimodal information across the brain to extend the flexibility for translating sensation into cognition, thereby facilitating advanced human mental and behavioral functions. Moreover, a recent study revealed a repeatable resting-state network partition based on the same Glasser-360 cortical parcellation used in the current study^37^. Resting-state networks related to primary functions (e.g., visual, somatomotor, and auditory) are dominantly enriched in a single type of shaker hub or stabilizer node, while the networks with high-order cognitive functions (e.g., default mode, frontoparietal, and cingulo-opercular) are heterogeneously depleted in more different dynamic types involving bi-active and switcher hubs. Together, the identified dynamic architecture reflects the functional division and hierarchy of large-scale functional networks in the human brain.

### Expression of ion channel and mitochondria-related genes underlies the intrinsic chronnectome

We found that the gene expression profiles that can explain the spatial heterogeneity in the dynamic characteristics of the chronnectome are mainly related to ion channels and mitochondria. Ion channels are crucial structures for the generation and conduction of electrical signaling in neurons that allow the movement of ions across the membrane, leading to changes in membrane potential and further neuronal electrical signals. The summed electric current flowing from multiple nearby neurons can generate local field potentials in the extracellular space around neurons, and the dynamic fluctuation of the local field potential is associated with the hemodynamics that can be captured by fMRI^38^. Thus, the dynamics of the macroscale brain networks have a tight association with the substantial neuronal electrical activity induced by ion channels. Our results further illustrate the molecular and biological foundations of the macroscale chronnectome in the resting human brain. In line with our results, previous findings from transcriptome-neuroimaging association studies have demonstrated that the expression of ion channel-related genes is closely related to static resting-state functional networks^18, 39^, dynamic streams in functional networks during tasks^21^, and structural networks^40–42^. Therefore, we speculate that the gene expression related to ion channels might be a transmodality and trans-state molecular basis for human brain networks.

The mitochondrion is the “powerhouse” organelle of the neuron and synthesizes adenosine triphosphate via the citric acid cycle and oxidative phosphorylation to provide energy for neuronal signaling and the maintenance of the resting membrane potential^43^. Our study revealed a significant coupling between the expression pattern of mitochondria-related genes and the intrinsic chronnectome pattern, with high expression dominantly located in the bi-active hubs (medial/lateral prefrontal and parietal cortices) that exhibit a high level of dynamic features. Consistently, the occupancy rate in the dynamic connectivity state linking most of these regions is associated with a single nucleotide polymorphism component with top related genes associated with metabolism^44^. These regions are essential to mediate various complex cognitive functions^45^ and are phylogenetically late developing, with a disproportionate enlargement during evolution^46^. Moreover, these regions exhibit high levels of energy metabolism, such as glucose utilization, oxygen consumption and regional cerebral blood flow^47, 48^. A recent study focusing on brain structure demonstrated that the variation of regional scaling to the normative brain size and shape in frontoparietal regions is associated with the expression of mitochondria-related genes^49^. Together with our results, these findings suggest that the large expansion and complexity in structures and functional dynamics are supported by substitutional energy production at the cellular level.

### Gene expression supports the regional involvement of the intrinsic chronnectome in behaviors

The architectures of the chronnectome during rest have been found to be associated with several specific aspects of cognitive performance, such as perception and executive function^10, 50^. Here, we demonstrated that intrinsic chronnectomes are involved in various cognitive and behavioral phenotypes. This association indicates a brain substrate for the well-known *g* factor proposed by Charles Spearman in the early 20th century, which represents a core mental ability shared among different cognitive and behavioral performances^51^. Thus, the chronnectome architecture likely contributes to the *g* factor to support a variety of cognitive and behavioral performances, which is consistent with the findings on the static connectome^28^. The differences in regional behavioral involvement between different types of dynamic nodes are mainly derived from their variances in modular reorganization, highlighting the great contributions of modular architectures to complex cognitions and behaviors^52, 53^.

We found that both the behavior involvements and the gene expression profiles varied among the four types of dynamic nodes. On the one hand, the diverse transcriptional signatures underlying the dynamic patterns suggest that the macroscale chronnectome architecture is largely shaped by micro-scale molecular mechanisms, such as neuroelectric activity and cellular metabolism. On the other hand, long-term involvement in functional processing may, in turn, alter the regional dynamic characteristics through a neuroplasticity mechanism^54^, further increasing the spontaneous dynamic heterogeneity across brain regions. These findings jointly suggest a complex interaction between gene regulation and function that shapes the spatial configurations of the intrinsic chronnectome in the human brain. To date, there have been extensive studies investigating gene-behavior, gene-brain and brain-behavior relationships in humans, which have provided plentiful valuable findings. Genes can indirectly modulate the behavioral phenotype by affecting brain structure and function via molecular and cellular mechanisms^55, 56^. Delineating the gene-brain-behavior pathways can expand our understanding of the linkage among these three aspects. Importantly, we found that regional behavioral involvement can be significantly predicted by the expression pattern of the chronnectome-related genes enriched for ion channels and mitochondria, potentially serving as a general mechanistic interpretation for a mediation router of micro-scale neuroelectric activity and cellular metabolism underlying brain-behavior coupling. Future studies simultaneously collecting gene expression, neuroimaging and behavioral data from large cohorts are urgently required for mediation analyses to assess gene-brain-behavior pathways at an individual level.

### Further consideration

First, the gene expression data from the AIBS were sampled from six donors with six left hemispheres and two right hemispheres. The limited sampling might have created a bias in capturing the variance in gene expression across individuals. Future studies with larger sample whole-brain genome-wide gene expression data could better address this issue. Second, the age of subjects used for rfMRI data and transcriptional data acquisition were not perfectly matched; the six brain samples from the AIBS (42.5 ± 13.4) were on average older than the 681 individuals from the HCP (28.7 ± 3.7). Given the significant age effect on both gene expression^57^ and functional brain architecture^58^, the difference in age could be a confounding effect. Third, we constructed the human chronnectome from multiband fMRI data. Although BOLD imaging captures hemodynamic signals that can indirectly reflect neural electrical activity^59, 60^, future studies with a simultaneous collection of fMRI and electrophysiological data would provide better insight into the understanding of gene-brain interactions. Finally, the findings reported here were correlational, not causative. Generally, we theoretically speculate that brain endophenotypes and behavioral phenotypes are largely regulated by gene expression. Determining whether and how the gene, brain, and behavior interact among one another remain interesting areas for further investigation.

## Methods

### HCP datasets

Data of 681 participants (ages 22-37 years, 381 females) with complete rfMRI data (two runs in each of the two sessions) from the S900 Data Release of HCP^23^ were analyzed. Ninety-two core cognitive and behavioral measures were selected from the datasets (*Table S1*). Multiband gradient-echo-planar whole-brain imaging acquisitions were acquired on a 3T Siemens Connectome Skyra scanner at Washington University, USA. The rfMRI data were obtained with the following sequence parameters: repetition time (TR) = 720 ms, echo time (TE) = 33.1 ms, flip angle (FA) = 52°, bandwidth = 2,290 Hz/pixel, field of view (FOV) = 208 × 180 mm^2^, matrix = 104 × 90, 72 slices, voxel size = 2 × 2 × 2 mm^3^, multiband acceleration factor = 8, and 1,200 volumes (14 min and 24 s) for each run. For each participant, four runs of rfMRI were scanned in two days, where one session containing two rfMRI runs was performed each day. During the scan, participants were instructed to keep their eyes open with a fixation. After the HCP minimal preprocessing procedure^61^, we further used SPM12 and GRETNA^62^ to reduce the biophysical noise in the rfMRI data by regressing out the linear trend, twenty-four head motion parameters, cerebrospinal fluid, white matter, and global signals and performing temporal bandpass filtering (0.01-0.1 Hz). For details, see *SI*.

### AIBS datasets

Brain gene expression data from six donors (mean age: 42.5 years, 1 female) were obtained from the AIBS^17^ (*Table S2).* Each postmortem brain was dissected into approximately 500 anatomically discrete samples per hemisphere. Briefly, each sample was spatially registered to the Montreal Neurological Institute (MNI) coordinate space according to the T1-weighted image obtained before dissection, and the locations of all samples are given in MNI coordinates in the SampleAnnot.csv file of each donor’s microarray data files. Normalization processes were conducted to minimize the potential effects of nonbiological biases and make the gene expression data comparable among the samples within and across brains. The detailed normalization methods are described in the technical paper for microarray data normalization (http://help.brain-map.org/display/humanbrain/Documentation?preview=/2818165/5177355/Normalization_WhitePaper.pdf). The normalized gene expression data of 58,692 probes were available for each sample.

### Constructing dynamic brain networks

We used a multimodal brain atlas^24^ to parcellate the cerebral cortex into 360 regions of interest. For each participant, the mean time series was extracted for each region based on the preprocessed rfMRI data. Dynamic functional networks were estimated by employing a widely used sliding window approach for each participant^4–6, 8–10^. Notably, the time window had a width of 100 s (i.e., 139 TRs) and a slide on time with a step of 0.72 s (i.e., 1 TR), allowing sufficient time points to estimate dynamic functional connectivity at the low-frequency band of interest (0.01-0.1 Hz) and simultaneously capturing temporal variations during a short period^6, 10, 63, 64^ Within each window, we estimated the functional connectivity matrix by calculating the Pearson’s correlation between any pair of network nodes based on the segments of the time series in the window. Therefore, for each participant, we obtained the dynamic networks for each of the four runs. Each of the dynamic networks had a total of 1,062 connectivity matrices (i.e., windows), and each matrix had 64,620 functional connections (i.e., 360 × 359/2). To validate the potential effect of the dynamic functional network construction approach on our main result, we also employed the framewise dynamic conditional correlation (DCC) method^25^ to reconstruct the dynamic networks and repeated our analysis. Please see “Validation of Effect of Dynamic Network Construction” for more details.

### Aligning AIBS datasets to brain atlas

To match the gene expression data from AIBS and the brain parcellation used in dynamic network construction, we performed several preprocesses for the gene expression microarray data of brain samples, including mapping samples to network nodes, probe reannotation and selection, and normalization across donors. Briefly, a total of 1,791 brain tissue samples from six donors were first referenced to the 360-region brain parcellation according to their MNI coordinates^18, 39, 40^. Specifically, if a sample did not fall within any region, we extended the matching range to a radius of 5 mm to search for the nearest brain region. The final distribution of the samples covered 84% of the parcellation (i.e., 301/360). Notably, the assigned samples were distributed evenly among the four categories of dynamic network nodes (*Table S4).* Subsequently, we used the Re-annotator toolkit^65^ to reannotate the gene assignment of probes with the reference genome assembly hg19^66^ and excluded probes that were not assigned to any gene and those with more than two mismatches between their sequences and reference. We also removed the probes without an Entrez ID or without significant calls in fewer than 50% of the assigned samples across all donors. These procedures resulted in 32,191 probes corresponding to 16,392 genes. Then, for each gene in each donor, we averaged the expression level across probes in each sample, followed by a Z-score normalization (subtracted the mean and divided by the standard deviation) across the samples and further averaging of the Z-scores of all samples within a network node. Finally, a gene expression map at the group level was obtained by averaging the Z-score of the gene expression level across the six donors.

### Identifying patterns of brain network dynamics and associating with cortical hierarchy

For each node, we computed two dynamic measures: the tGV, which represents the overall fluctuation amplitude of its connections, and the tMV, which represents the degree of variation degree of module affiliations across time^6^, where the module affiliation was determined using the InfoMap algorithm^67^ (see *SI* for details). Based on the group-level, session-averaged tGV and tMV maps, all brain nodes were assigned into four distinct categories according to their comparisons with the mean value of each measure. We also repeated this classification analysis in each session and calculated normalized mutual information (NMI) between the node classification pattern of the two sessions to estimate the between-session reliability (Fig. S1A and Fig. S1B). To further illustrate the basic dynamic information of different dynamic hubs, we calculated the power spectrum of overall connection dynamics and modular switching frequency for each node and compared these characteristics across the four node-categories. Moreover, we performed two asymmetry analyses to delineate the inter-hemispheric similarity of the tGV and tMV maps (see SI).

We then explored the consistency between the spatial layout of network dynamics and cortical hierarchy^11^ by calculating the percentiles of each category of the network nodes within each type of hierarchy area. Permutation tests were utilized to determine whether the percentiles were larger than those by chance. Moreover, we utilized a publicly available myelin map derived from T1- and T2-weighted MRI^14, 24^ to study the myeloarchitectural feature of the dynamic network patterns. The myelin contents were allocated for each node and were compared between each category pair using permutation tests.

### Association between brain network dynamics and transcriptional signatures

To explore the molecule mechanism underlying the spatial layout of intrinsic network dynamics, we utilized a partial least squares regression model^40, 68^ in which the gene expression data of brain regions (301 nodes × 16,392 genes) were set as the predictor variables and the tGV and tMV of brain regions (301 nodes × 2 measures) were set as the response variables (*SI).* A permutation test was performed to determine the statistical significance of each component (10,000 times). Then, for each significant component, we used a bootstrapping method to assess the estimation error of the weight for each gene and further divided the weight by the estimated error to obtain the corrected weight for each gene^40^. We ranked the genes according to their corrected weight, which represents their contribution to the partial least squares regression component. Finally, the Gene Ontology enrichment analysis and visualization tool (GOrilla, http://cbl-gorilla.cs.technion.ac.il/)^69, 70^ was used to identify the enriched Gene Ontology terms of the ranked genes from each significant component.

To further validate the robustness of the results in the partial least squares regression and enrichment analysis, we also employed an internal replication strategy. Briefly, we split the rfMRI data of the HCP datasets into two cohorts with matched age, gender and handedness and obtained the group-level tGV and tMV maps within each subgroup (i.e., HCP-cohort1 and HCP-cohort2). For the gene expression data from the AIBS datasets, we split six brains into two subgroups as in a previous study^18^, in which each subgroup included a full brain (H0351.2001 or H0351.2002) and two half brains from the other donors to balance the sample size, resulting in six possible subgroup pairs (see *Table S6*). For each subgroup pair of AIBS datasets, we repeated the partial least squares regression with two cohorts of HCP (i.e., 2 subgroups × 6 pairs × 2 cohorts = 24 repetitions). Then, we performed the enrichment analysis for each repetition and compared the results with those obtained in the main analysis.

### Association between transcriptional signatures and behavioral involvement of brain dynamics

An epsilon-insensitive support vector regression with radial basis function kernel was used to predict the behavioral involvement of brain regions. A leave-one-region-out strategy was used for cross-validation. The expression levels of top-ranked genes (top 10%) in two significant components from partial least squares regression were concatenated to produce a predictive feature vector. To evaluate the behavioral involvement of node dynamics, we performed a canonical correlation analysis (CCA) with two dynamic measurements (the tGV and the tMV) and ninety-two cognitive and behavioral measurements as two variable sets. The behavioral involvement for each region was calculated by summing the total absolute weight in the significant CCA mode on two dynamic measurements. Finally, a permutation test^71^ was performed to estimate the statistical significance of the observed prediction accuracy of the support vector regression (10,000 times).

### Validation of the effect of head motion

We excluded participants with large head motion (above 3 mm or 3° in any direction and in any run) and regressed out individual mean framewise displacement in the canonical correlation analysis. To further strictly control the potential influence of head motion on our main findings, we performed a spike regression-based scrubbing in the nuisance regression procedure^29, 30^ during preprocessing with the criterion of a framewise displacement above 0.5 mm and repeated our main analyses above.

### Validation of the effect of dynamic network construction

To validate the potential effect of the sliding window-based network construction approach on our main results, we also employed the framewise dynamic conditional correlation (DCC) method^25, 72^ to reconstruct the dynamic networks and repeated our analysis. The DCC calculates inter-regional dynamic correlations by estimating the potentially large covariance structures with the sequential estimation scheme and extremely stingy parameterization. Briefly, the DCC algorithm mainly includes two steps. First, a univariate generalized autoregressive conditional heteroskedasticity (GARCH) model was fit to each time course and further used to estimate standardized residuals. Second, an exponential weighted moving average (EWMA)-type approach was applied to these standardized residuals to compute the time-varying correlation. Please refer to Lindquist *et al*.^25^ for details on the DCC computations.

## Acknowledgments

The authors would like to thank Dr. Yanchao Bi and Dr. Tengda Zhao for helpful discussions. This work was supported by the National Natural Science Foundation of China (81620108016, 31830034, 81671767), the Changjiang Scholar Professorship Award (T2015027), and the Fundamental Research Funds for the Central Universities (2017XTCX04). Imaging data were provided by the Human Connectome Project, WU-Minn Consortium (Principal Investigators: David Van Essen and Kamil Ugurbil; 1U54MH091657) funded by the 16 NIH Institutes and Centers that support the NIH Blueprint for Neuroscience Research; and by the McDonnell Center for Systems Neuroscience at Washington University. The authors thank the Allen Institute for Brain Science.

## Author contributions

Y.H., J.L. and M.X. conceived the study; J.L. and M.X. performed the data analysis with technical support from X.W. and X.L.; J.L., M.X. and Y.H. wrote the manuscript; all authors commented on the study and manuscript.

